## Supplementary material for "Multi-parameter photon-by-photon hidden Markov modeling": SI

[1]Department of Biological Chemistry, The Alexander Silberman Institute of Life Sciences, Faculty of Mathematics & Science, The Edmond J. Safra Campus, The Hebrew University of Jerusalem, Jerusalem 9190401, Israel
[2]Physical and Synthetic Biology. Faculty of Biology, Ludwig-Maximilians-Universität München, Großhadernerstr. 2-4, 82152 Planegg-Martinsried, Germany
[3]Department of Chemistry and Biochemistry, and Department of Physiology, University of California, Los Angeles, California, United States
[4]Department of Physiology, CaliforniaNanoSystems Institute, University of California, Los Angeles, California, United States
[5]The Center for Nanoscience and Nanotechnology, The Hebrew University of Jerusalem, Jerusalem 9190401, Israel

October 13, 2021

#### S1 Data Analysis

##### S1.1 H$^2$MM analysis

All code and raw data (in the form of photon-HDF5 files) is available for download here: ( https://zenodo.org/record/4671393 [1]). The repository contains .yml files to directly recreate the Anaconda environments. On Windows platforms make sure in install Visual Studios (we have tested 2015 and 2019) with all options installed. FRETBursts and H2MM_C may exhibit difficulties in

*
†

installation. In that case, please install these repositories with the following commands from your terminal:

- H2MM_C is always installed with:

```
pip install git+https://github.com/harripd/H2MMpythonlib.git
```

- FRETBurts should be installed with one of the following:
  - For the mph2mmenv.yml environment use (this is the package used by all notebooks except the anisotropy notebook):

    ```
    pip install git+https://github.com/harripd/FRETBursts.git
    ```

  - For the mph2mmpolenv.yml environment use (this is only for the anisotropy notebook):

    ```
    pip install git+https://github.com/harripd/FRETBursts.git@polarization
    ```

  - For the oldversion.yml use (this is for those using Python 3.7, you will have to fix the sns.kdeplot function calls to fit the older signature):

    ```
    conda install fretbursts -c conda-forge
    ```

Data must be extracted from the FRETbursts[2] data structure, and cast such that the H2MM_C package[3] can process it. In the $H^2$MM algorithm, photons are identified by an unsigned integer index, and an unsigned integer arrival time. A burst consists of two arrays of equal length, one for the photon indexes, and the other for the photon arrival or detection times (in the form of one dimensional numpy arrays). The python function accepts as input an initiating $H^2$MM state model (implemented as a python extension type in the H2MM_C package[3]), a python list of the arrays of the photon indexes, and a separate python list of the arrays of the photon arrival times.
FRETbursts identifies streams according to the following convention: the excitation period is identified as either $D_{ex}$ or $A_{ex}$ for photons originating from donor or acceptor excitation, respectively. Similarly, the detector at which the photon arrived is identified by either $D_{em}$ or $A_{em}$ for donor or acceptor emission, respectively. Thus, as an example, a photon originating from donor excitation and arriving at the acceptor detector will be in the $D_{ex}A_{em}$ stream. In sp$H^2$MM, only the $D_{ex}D_{em}$ and $D_{ex}A_{em}$ photon streams are used, with the $D_{ex}D_{em}$ photon stream assigned index 0, and the $D_{ex}D_{em}$ photon stream is assigned index 1. For mp$H_2$MM, the same convention is used, but the $A_{ex}A_{em}$ photon stream is also included, and assigned index 2. The $A_{ex}D_{em}$ photon stream should be exclusively background, and is therefore discarded for mp$H^2$MM analysis, however, it should be noted that the H2MM_C package is fully capable of incorporating the $A_{ex}D_{em}$, or other photon streams, the exclusion is purely because the

$A_{ex}D_{em}$ photon stream should not contain any useful information in alternating laser excitation (ALEX)[4, 5] experiments. Generally, any photon stream that contains useful information should be included in the analysis. Optimizations are run for a maximum of 3,600 or 7,200 iterations, or until the improvement in the log-likelihood values between state-models is less than $10^{-7}$, at which point the improvement between iteration is insignificant. After model optimization, the *Viterbi* algorithm is used to both find the most likely state path through the data and to calculate the ICL[6, 7].
For all data sets, $H^2$MM state models are optimized with increasing numbers of states, until all of the following requirements were met: (i) a minimum ICL is found, (ii) a model with BIC' less than 0.005 is found (see section S2). We always start from the one state model to serve as a baseline, and up to at least one more state that is required by prior knowledge of the system and visual inspection of the E-S plots. We always fit a minimum of four states, because visual inspection of the E-S plots and BVA plots of all of our samples seem to have at least two FRET states, a dark donor state, and a dark acceptor state. The function implementing the *Viterbi* algorithm in H2MM_C also sorts and characterizes the dwells automatically.

###### S1.1.1 The $A_{ex}$ shift procedure in µsALEX

The $A_{ex}A_{em}$ time shift is used in µsALEX data, and is built into the function that converts FRETBursts data into a format readable by H2MM_C. There are three types of time shifts that could be employed: a simple *shift*, a *random* shift, and finally an *even* option. The basic principle is to shift $A_{ex}A_{em}$ photons into the time window of donor excitation, while still applying the smallest change in the arrival time. In principle this means pairing donor and acceptor excitation periods into adjacent periods and reassigning each $A_{ex}A_{em}$ photon into its paired period. The *shift* option simply takes the difference between the beginnings of donor and acceptor excitation periods, and subtracts that value from all $A_{ex}A_{em}$ photon arrival times. The *random* option assigns random times to each $A_{ex}A_{em}$ photon within its adjacent donor excitation period. The *even* option redistributes all $A_{ex}A_{em}$ photons in a given acceptor excitation period into the adjacent donor excitation period, such they they are equidistant in time from one another. In the YopO data, the *even* option is always used.
Assesment of µsALEX data should use the *even* option for application of the $A_{ex}$ shift. The *random* option is generally advisable only as a control for results fit with the *even* option.

###### S1.1.2 µsALEX cautions

The shift of $A_{ex}A_{em}$ photons presents significant potential for artefacts, and should always be used cautiously. Because photons are redistributed, a portion of information from the arrival times of the $A_{ex}A_{em}$ photons is lost. This also produces a series of gaps in photon arrival times in the shifted data, as no photons arrive during the $A_{ex}$ period. Changes in the $A_{ex}A_{em}$ stream that

occur at timescales similar to the alternation period are essentially erased with the shift.
Further research is warranted on better ways of enabling analysis of µsALEX data. These could come in two forms:

1. Advanced shifting procedures, which maintain more information about the arrival times, potentially through modeling of the $A_{ex}A_{em}$ stream.

2. Modification of the $H^2$MM algorithm to penalize transitions with overly regular transitions that do not distribute exponentially.

**Rapid dynamics in µsALEX** The YopO data presented in the main text provides a test case to consider these limitations. The total duration of an entire $D_{ex}$ and $A_{ex}$ cycle is 50 µs, which gives an alternation rate of 20 kHz. Therefore, if any transition rate constants that are similar to 20,000 $s^{-1}$ are encountered, suspicion is warranted. In the YopO data, the fastest transition rate constant is 13,000 $s^{-1}$, which is the transition rate from the high FRET state to the low FRET state, in the apo form. The reverse process is slower, with a rate constant of 6,000 $s^{-1}$. These rates are within an order of magnitude of the alternation rate, and therefore closer examination is necessary to determine whether or not these are artefacts or represent real transition rates. The first observation is that the two states involved in these transitions are both FRET states, and have similar $S_{raw}$ values. Since this transition does not rely heavily on the $A_{ex}A_{em}$ stream, which is where artefacts might arise, the transition rates may be accurate. However, the typical dwell in these states will be similar to the alternation period. This may mean that many $A_{ex}A_{em}$ photons will have been redistributed between transitions between the high and low FRET states, resulting in unreliable $S_{raw}$ values.
To address these concerns, we repeat the YopO experiments under identical conditions except for changing the alternation period. We try 20 and 100 µs alternation periods. In the 20 µs alternation period data, the laser ramping of the donor laser resulted in an unfeasibly short window of donor excitation for the data to be usable. The 100 µs alternation period data, however, is usable. This alternation period corresponds to an alternation frequency of 10 kHz, which is very close to the transition rate that we extract. When mp$H^2$MM is applied to this data, the equivalent transition rate is 10,000 $s^{-1}$. As this is identical to the alternation rate, it is therefore suspicious as an artefact. This is expected given the 50 µs alternation period data, as it predicts a rate faster than but similar to the 100 µs alternation period, and therefore in the region where artefacts are likely.
As another test, we use dwell-time analysis on results from the *Viterbi* algorithm. Real processes should result in exponentially distributed dwell times, while if the models contain artefacts, we would expect the distribution to be heavily weighted towards dwells with durations around one or a small number of alternation periods. For the data with both 50 and 100 µs alternation periods, the dwell times in the low FRET state distribute exponentially, while

those in the high FRET state appear to be bi-exponential (see supplementary figures S26,S27). The dominance of the longer lifetime, however, indicates that an influence of the alternation period, if any, is minimal. This serves as a strong evidence that the 50 µs alternation period data is reliable. The dwell times also indicate that the 100 $s^{-1}$ alternation period data, while similar to the alternation rate, may not be an artefact.
We also try two additional methods to test if the rate constant values we retrieve are a result of an artefact or not. The first is to perform analyses of randomly chosen sub samples of the data (see supplementary section S1.4 for implementation details). This allows us to take a the value of a given transition rate constant, and test how much the optimized values vary depending on the sub sample. The other is to take sub samples from the 50 and 100 µs alternation period data of the same system under measurement, combine the subsets, and perform mp$H^2$MM fitting on these combined sets. Using these two approaches, we assess the statistical uncertainty in the estimation of the rate constant values, from two different alternation periods, and test whether the differences in the rate constant estimates are within the range of statistical uncertainty. Using these approaches similar standard deviations of the state model parameters are retrieved, regardless of the alternation periods of the measurement (see Jupyter notebook YopO_usALEX_error_and_artefact_testing in supplementary data set [1]). Results are similar when combining bursts from both alternation periods. Therefore, we conclude that the rate constants retrieved from analyzing the 50 µs alternation period data is not an artefact.

**Further discussion** Our 50 µs alternation µsALEX data proved reliable, while the 100 µs data potentially only requires longer acquisition. Now, we turn to discuss the potential pitfalls, where values should not be considered reliable. Suppose a system where the model exhibits or retrieves transition rates between two states with drastically different $S_{raw}$ values that are similar to the alternation period. In this case, because the two states are distinguished more so by the $A_{ex}A_{em}$ stream, such a model should not be considered reliable in any way, as those states are very likely the result of an artefact resulting from the shift procedure used in the treatment of µsALEX data within mp$H^2$MM. Transition rates that are very similar to small integer multiples or fractions of the alternation period should also be considered suspicious. For instance, given a 50 µs alternation period, transition rates that are close to 10,000 $s^{-1}$ are more likely to be an artefact than the 13,000 $s^{-1}$ transition rate seen in the YopO data.
Whenever transition rates are within an order of magnitude of the alternation rate, especially ones that are small integer multiples or fractions, further analysis will be needed to confirm that the extracted rates and states are not artefacts. If the transition is between two states of similar $S_{raw}$, then the rates and $E_{raw}$ values are more likely to be reliable, but the $S_{raw}$ values should be considered to be strongly influenced by the other state. If, however, the transition is between two states with significantly different $S_{raw}$, then it is likely to be an artefact,

especially if the reverse rate is also similar to the alternation rate. Any time either scenario is encountered, it should be considered mandatory to perform additional analysis. We present three primary methods to conduct this:

1. Dwell time analysis using the *Viterbi* algorithm: dwell times should distribute exponentially, with no sudden drops in the histogram.

2. Analysis of variance of subsets.

3. Measure with a different alternation rate: transition rates and $E_{raw}$ values should be similar within the recovered statistical uncertainty.

Tests of dwell time analysis are recommended at all times to serve as a general check for any potential problems with the data. Users should always be sure to examine these models closely, and take all prior knowledge into account when analyzing this data. In summary, the shift is a work-around for µsALEX experiments, and great care and examination should be employed in using it. Ultimately, nsALEX should always be preferred for mpH$^2$MM analysis, and when µsALEX is used, the limitations of must be understood.

#### S1.2 Parameter Calculations

The transition probability matrix contains the values of the transition rate constants in inverse units of the clock period of the measurement, that is the time period corresponding to one time interval of the measurement. For these measurements this is the pulsed laser repetition rate, which is 50 MHz in our experiments. Calculation of $E_{raw}$ and $S_{raw}$ is slightly more complicated, especially for mpH$^2$MM. Following the nomenclature of Lee *et al.*[8], $E_{raw}$ is generally defined as in Eq.S1:

$$E_{raw} = \frac{F_{D_{ex}}^{A_{em}}}{F_{D_{ex}}^{A_{em}} + F_{D_{ex}}^{D_{em}}} \tag{S1}$$

where $F_{D_{ex}}^{A_{em}}$ indicates the raw, uncorrected counts of acceptor photons during donor excitation period, (the counts in the $D_{ex}A_{em}$ photon stream) and $F_{D_{ex}}^{D_{em}}$ is likewise the raw, uncorrected donor counts during donor excitation period (the counts in the $D_{ex}D_{em}$ photon stream). $S_{raw}$ is likewise defined in Eq.S2:

$$S_{raw} = \frac{F_{D_{ex}}^{A_{em}} + F_{D_{ex}}^{D_{em}}}{F_{D_{ex}}^{A_{em}} F_{D_{ex}}^{D_{em}} + F_{A_{ex}}^{A_{em}}} \tag{S2}$$

where, following the nomenclature $F_{A_{ex}}^{A_{em}}$ is the raw, uncorrected acceptor emission during acceptor excitation period (the counts in the $A_{ex}A_{em}$ photon stream). These values have clear equivalents in the emission probability matrix. From Pirchi *et al.*[9] the emission probability matrix is defined as in Eq. S3:

$$(\hat{B}_{i,k}) \equiv b_{i,k} = P(y_t = k | x_t = i, \hat{\lambda}),\ for\ all\ t \quad and \begin{Bmatrix} 1 \leq i \leq N_s \\ 1 \leq k \leq N_p \end{Bmatrix} \tag{S3}$$

where $\hat{B}_{i,k}$ is the emission probability matrix, $i$ is a given state, $k$ is a given photon stream, $y_t$ is the photon at time $t$, $x_t$ is the state the system is in at time $t$, and $\hat{\lambda}$ is the $H^2MM$ model. $N_s$ and $N_p$ are the maximum number of states and photon streams, respectively.
It is also required that the emission probability matrix be row stochastic, as defined in Eq. S4:

$$\sum_{k=1}^{N-s} b_{i,k} = 1 \tag{S4}$$

Practically, we view the emission probability matrix as being the probability that a photon will belong to the stream $k$ given the molecule is in state $i$. Therefore, we can make an equivalence between the $F$ values from Lee *et al.* and the elements of $b$ as in Eq. S5:

$$_iF_{k_{ex}}^{k_{em}} \equiv b_{i,k} \tag{S5}$$

Therefore, ${}^iE_{raw}$ and ${}^iS_{raw}$ of each state in a given $H^2MM$ model are defined as in Eq. S6 and S7:

$${}^iE_{raw} = \frac{b_{i,DexAem}}{b_{i,DexAem} + b_{i,DexDem}} \tag{S6}$$

and

$${}^iS_{raw} = \frac{b_{i,DexAem} + b_{i,DexDem}}{b_{i,DexAem} + b_{i,DexDem} + b_{i,AexAem}} \tag{S7}$$

It should be noted that given the requirement of row stochasticity, the denominator of an $spH^2MM$ model will be 1 in Eq. S6, and therefore the ${}^iE_{raw,spH^2MM} = b_{i,DexAem}$, but this does not hold true for $mpH^2MM$. Therefore practitioners of $spH_2MM$ may be accustomed to simply looking at $b_{i,DexAem}$ as the ${}^iE_{raw}$, and must be careful to discontinue this practice when moving to $mpH^2MM$.

#### S1.3 Viterbi Analysis

The $H^2MM$ algorithm has no direct means to assess the error on individual values within a given state model for a given data set. However, the *Viterbi* algorithm provides a convenient way of obtaining a proxy for the error of individual values in a given $H^2MM$ model. The process is the same as in Lerner *et al.*[10]. After *Viterbi* analysis, consecutive photons classified as belonging to the same state are grouped into dwells.

##### S1.3.1 $E_{raw}$ and $S_{raw}$ values

The counts of photons in each stream can then be used to assign each dwell a mean $E_{raw}$ value, and for $mpH^2MM$, also a mean $S_{raw}$ value. The definitions are essentially the same as those for bursts from Lee *et al.*[8], and are given in Eq. S8 and S9:

$$E_{raw,dwell} = \frac{n_{dwell}^{DA}}{n_{dwell}^{DA} + n_{dwell}^{DD}} \tag{S8}$$

$$S_{raw,dwell} = \frac{n_{dwell}^{DA} + n_{dwell}^{DD}}{n_{dwell}^{DA} + n_{dwell}^{DD} + n_{dwell}^{AA}} = \frac{n_{dwell}^{DA} + n_{dwell}^{DD}}{n_{dwell}^{total}} \tag{S9}$$

where $n_{dwell}^{stream}$ is the number of photons in originating from the given photon *stream*, with the first letter denoting the excitation period, and the second the detection channel. It should be noted that in spH$^2$MM $n_i^{DD} + n_i^{DA} = n_i^{total}$, and $S_{raw,dwell}$ cannot be calculated. It is then possible to define a ${}^S\bar{E}_{raw,w}$ and ${}^S\bar{S}_{raw,w}$ for each state $S$. As the total number of photons in each dwell varies, which could bias a traditional mean, we opt to use a weighted average instead, as defined in SI Eq S10 and S11:

$$^S\bar{E}_{raw,w} = \frac{\sum_i^S (n_i^{DD} + n_i^{DA}) E_{raw,i}}{\sum_i^S (n_i^{DD} + n_i^{DA})} \tag{S10}$$

$$^S\bar{S}_{raw,w} = \frac{\sum_i^S n_i^{total} S_{raw,i}}{\sum_i^S n_i^{total}} \tag{S11}$$

where $n_{dwell}^{total}$ is total number of photons in the dwell in all streams.
Their standard deviations (SD), as defined in SI Eq S12 and S13:

$$SD(^S\bar{E}_{raw,w}) = \left[\frac{\sum_i^S (n_i^{DA} + n_i^{DD})(E_{raw,i} - {}^S\bar{E}_{raw,w})^2}{\sum_i^S (n_i^{DA} + n_i^{DD})}\right]^{1/2} \tag{S12}$$

$$SD(^S\bar{S}_{raw,w}) = \left[\frac{\sum_i^S n_i^{total}(S_{raw,i} - {}^S\bar{S}_{raw,w})^2}{\sum_i^S n_i^{total}}\right]^{1/2} \tag{S13}$$

where $l_S$ is the number of dwells in state $S$.
Their standard errors (SE), as defined in SI Eq S14 and S15:

$$SE(^S\bar{E}_{raw,w}) = \left[\frac{\sum_i^S (n_i^{DA} + n_i^{DD})(E_{raw,i} - {}^S\bar{E}_{raw,w})^2}{\sum_i^S (n_i^{DA} + n_i^{DD})}\right]^{1/2} / \sqrt{l_S} \tag{S14}$$

$$SE(^S\bar{S}_{raw,w}) = \left[\frac{\sum_i^S n_i^{total}(S_{raw,i} - {}^S\bar{S}_{raw,w})^2}{\sum_i^S n_i^{total}}\right]^{1/2} / \sqrt{l_S} \tag{S15}$$

##### S1.3.2 Dwell time analysis

The difference in photon arrival times between the first photon in the dwell and the photon after the final photon in a dwell is characterized as the dwell duration. This holds true unless the dwell is the final one in the burst, in which case the difference is between the first and final photon in the dwell. A mean dwell time of dwells beginning in state $g$ and transitioning to state $h$ is then defined as in Eq. S16:

$$\bar{t}_{g,h} = \sum_{i=1}^{l_{g,h}} t_{i,g,h}/l_{g,h} \tag{S16}$$

The transition rate is $k_{g,h} = 1/\bar{t}_{g,h}$. The standard error of $t_{g,h}$ is defined as in Eq. S17:

$$SE(\bar{t}_{g,h}) = \left[ \sum_{i=1}^{l_{g,h}} (t_{i,g,h} - t_{dw\bar{e}ll,g,h})^2/l_{g,h}] \right]^{1/2} / \sqrt{l_{g,h}} \tag{S17}$$

where $l_{g,h}$ is the number of dwells that start in state $g$ and transition to state $h$. With the SE of the transition rate begin defined as in Eq. S18:

$$SE(k_{g,h}) = \frac{SE(\bar{t}_{g,h})}{t_{\bar{g},h}} \tag{S18}$$

Occasionally, the *Viterbi* algorithm predicts dwells with a very small number of photons. As these are likely spurious, we first exclude dwells from the analysis with five photons or less. Dwells at the beginning and end of bursts are truncated, and therefore we generally exclude them from the analysis to prevent them from biasing the results. However, when the transition rates approach the time scales of the burst durations (in the millisecond range), few dwells have a mean dwell time in the burst, making the analysis often misleading. Therefore, we choose to analyze the durations of the dwells at the beginnings and ends of bursts as separate data sets. Differences in these three data sets and comparison with the mean dwell time must all be taken into consideration in assessing the validity of the extracted transition rates. On the whole, however, we find that the transition probability matrix usually provides more reliable values than *Viterbi* derived mean dwell times. We also use the *Viterbi* results to flag transition rates as potentially spurious that have fewer than ten detected dwells that have more than five photons in them.

##### S1.3.3 Optional: Analysis of Photon Nanotimes

This work did not discuss fluorescence lifetime data. Nevertheless, fluorescence lifetime data can be derived from nsALEX experiments. Therefore, we hereby describe how to extract it and analyze it, if relevant. As the photon arrival or detection time relative to the last laser pulse (the photon nanotime) is also recorded, the fluorescence lifetime of bursts and sub-populations can be assessed. Using the *Viterbi* algorithm, each photon is assigned a sub-population.

A histogram of photon nanotimes of all photons is then assigned to the same sub-population and stream, resulting in separate nanotime histograms, or better put fluorescence decays, for each photon stream in each sub-population. Since the *Viterbi* algorithm also includes a posterior probability for each photon,providing a likelihood that the assignment is correct, we implement a threshold, removing photons with a posterior probability of less than 0.2 from consideration. Doing so, we find that changing the threshold does not have a significant impact on the shape of the histograms, and thus the threshold is somewhat arbitrarily set, as a trade-off between number of photons in the histogram reducing noise, and confidence of each photon assignment. See section S3 for a method to transform photon nanotimes into a parameter that can be characterized by the mpH$^2$MM approach.

##### S1.4 Error Analysis using sub samples

To asses parameter errors, we implement a function that performs a sequence of H$^2$MM optimizations on $N$ sub samples of the data given as input. Each sub sample is constructed to be of equal size, and no bursts are repeated between subsets. The first sub sample is composed of the bursts with indexes $1, N, 2N, 3N...$, the second of indexes $2, N+1, 2N+1, 3N+1$. The following sub samples all follow the same formula. Then, a list of state-model parameter values is constructed and serves as the basis for testing the their statistical uncertainty. The standard deviation of each parameter value of the models is taken, and used as the error for that value. It is important to mention that in freely-diffusing confocal-based smFRET, the majority of single-molecule photon bursts describe different un-synchronized molecules out of the bulk. Therefore, it is safe to estimate the state-model of a randomly chosen sub sample of bursts, as long as i) the sub samples include a considerably large amount of bursts, ii) that many sub samples are tested, and iii) that the overlap between the sub samples is minimized, or even non-existent, as in our case.

#### S2 Selection of a State-Model

While the minimum ICL is usually the best predictor of the ideal state-model, it is not perfect. Selection of an ideal model must take all other factors into account, primarily the BIC', BVA signatures of within-burst dynamics, and prior knowledge of the system. As noted in the main text the ICL is the most reliable means to rely on when selecting a most likely state model. However, we never relied exclusively on the ICL. We have yet to see a counter example to the number of states predicted by assessment of the ICL to be greater than the number of states predicted by assessment of BIC'. Therefore, we consider the ICL to be more conservative than BIC', and we usually find that the ICL provides more reasonable models. But, when the ICL- and BIC'-based model selections disagree, we consider both, especially when the disagreement is by two or more states. *Viterbi* analysis can also be useful, to see if dwells have

${}^{S}E_{raw,w}$ and ${}^{S}S_{raw,w}$ values that are centrally distributed about the $E_{raw}$ and $S_{raw}$ values predicted by the model, if they are not, it is reason to be suspicious of the model.

### S3 Transforming a non-binomially distributed variables for work with mpH$^2$MM

Consider the variable $t$, which is best described as distributing as $\mathbf{P}(t)$. We want to map it to a parameter $t'$ distributed by the beta distribution $\mathbf{B}(t')$. Since both are probability density functions (PDFs), their mapping should be performed on a basis of equal probabilities, which can be assessed according to their corresponding cumulative distribution functions (CDFs):

$$p = \int_{t_{min}}^{t} \mathbf{P}(t)\, dt = \int_{t_{min}}^{t} \beta(t')\, dt \tag{S19}$$

Eq. S19 will be the basis for transformation of a given $t$ value to a given $t'$ value. After this transformation, each $t$ value can be transformed into a $t'$ value for use as a parameter in the framework of mpH$^2$MM.
Let's take for example the donor photon nanotimes of in smFRET measurements with acquired photon nanotimes, such as nsALEX. We can accumulate all donor photon nanotimes into a histogram, better known as a fluorescence decay. Then, we can fit it with a sum of a few exponential functions, and then define $\mathbf{P}(t)$. We would then need to define the exact $\alpha$ and $\beta$ parameter values of the Beta distribution $B(t')$, to which we would like to map the photon nanotime data. A parameter continuous scale should be decided. We presume the scale should be mean lifetime going from 0 ns and all the way to the intrinsic dye lifetime, in the absence of acceptor, $\tau_D$. This should then be compared with the scale of mean values in the Beta distribution, from 1 to 0, respectively:

$$\langle\tau\rangle \in \{0, \tau_D\} \Rightarrow \langle t'\rangle \in \{1, 0\} \tag{S20}$$

Then, we can use Eq. S19 to start mapping photon nanotime of a given photon stream to a parameter that will correspond to the framework of mpH$^2$MM.

### S4 System Requirements

All notebooks in the Zenodo repository[3] were performed on an HP Pavilion Gaming 15-0001nx Laptop
CPU: Ryzen 7 3750H (4 cores, 2.3 GHz 6 MB cache) RAM: 16 GB DDR4-2400 SDRAM (2x8GB) Graphics Card: NVIDIA GeForce GTX 1660 Ti (6 GB GDDR6 dedicated) Operating system: Manjaro Linux, kernel Linux 5.10.63-1-MANJARO(x86_64)
We have also tested it with Intel i7 processors running Ubuntu, and Windows 10 systems. We recommend using at least an Intel i5 or equivalent. We have

not tested this on a system with less than 16GB of RAM, but based on memory usage, smaller datasets like HP3 should be able to run with 8GB, while larger sets like those used in MalE and YopO will probably require 16GB. H2MM_C does not use the graphics card, so graphics requirements will be determined by your operating system.

### S5 Supplementary Data

#### S5.1 Figures

##### S5.1.1 Expanded Introductory Figure

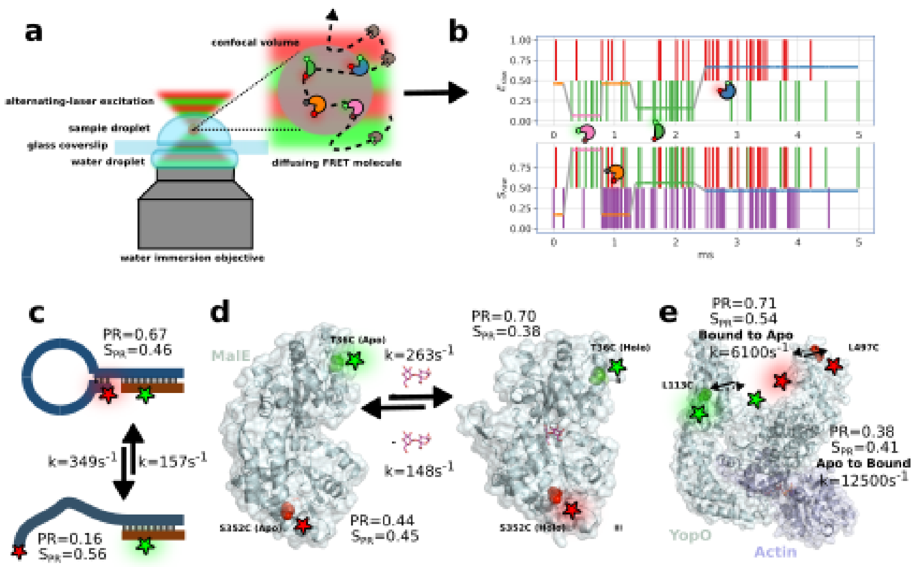

**figure** S1: **Cartoon representations of data acquisition, and biological systems examined in this paper** a) Confocal microscope setup with inset showing single molecules diffusing in and out of the confocal volume undergoing conformational and photophysical changes, producing b) a photon time trace, photons represented by vertical bars, and the most likely state-path according to the *Viterbi* algorithm overlayed as horizontal colored line. c-e) biological systems studied: c) DNA hairpin, d) maltose binding protein MalE conformational changes, and e) type III secretion effector YopO, along with mpH$^2$MM derived $E_{raw}$, and $S_{raw}$ values and the accompanying transition rates are give for those displaying exclusively intrinsic dynamics.

##### S5.1.2 Simulations

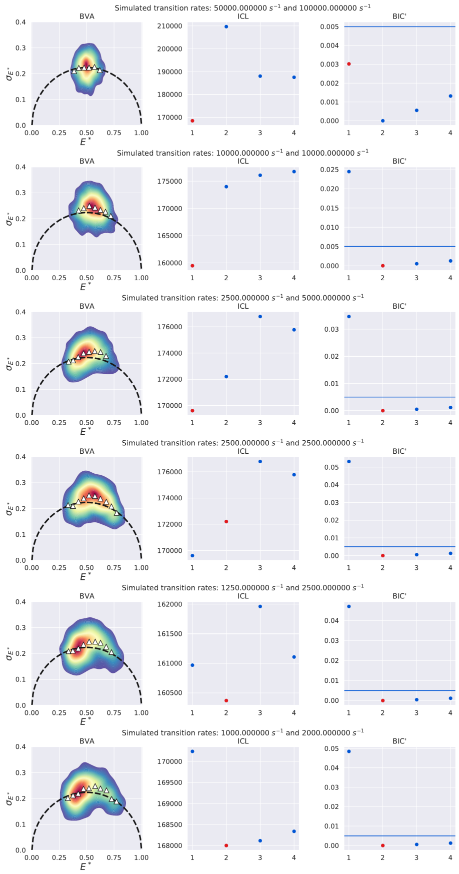

**figure** S2: **Simulations testing reliability of ICL.** Simulations conducted using PyBroMo with different transition rates, and the $ICL$ and $BIC'$ values of $H^2$MM fits. Note how ICL is minimized for slower transition rates.

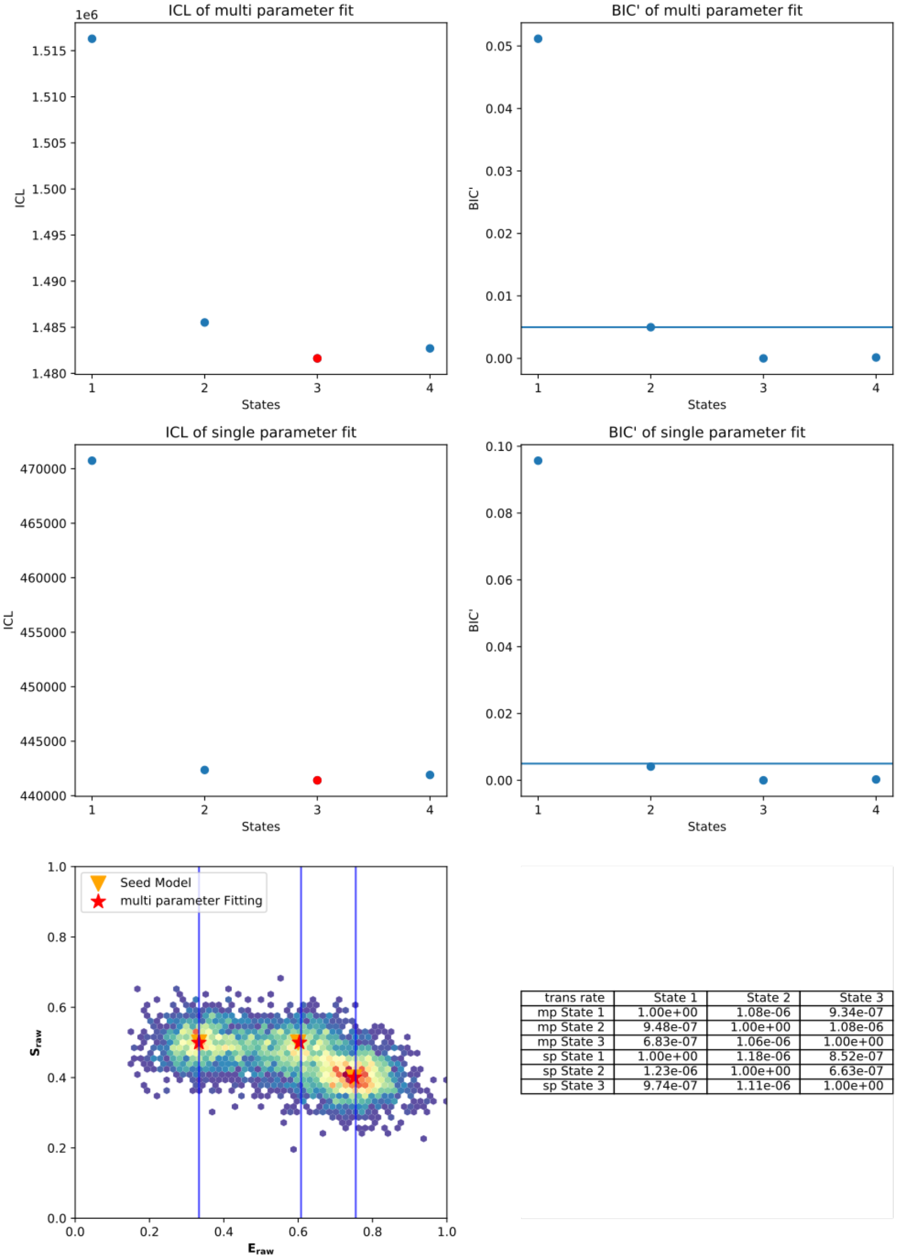

| trans rate | State 1 | State 2 | State 3 |
|---|---|---|---|
| mp State 1 | 1.00e+00 | 1.08e-06 | 9.34e-07 |
| mp State 2 | 9.48e-07 | 1.00e+00 | 1.08e-06 |
| mp State 3 | 6.83e-07 | 1.06e-06 | 1.00e+00 |
| sp State 1 | 1.00e+00 | 1.18e-06 | 8.52e-07 |
| sp State 2 | 1.23e-06 | 1.00e+00 | 6.63e-07 |
| sp State 3 | 9.74e-07 | 1.11e-06 | 1.00e+00 |

**figure** S3: **Simulation of 3 states with similar $\mathbf{S_{raw}}$** Both sp and mpH$^2$MM are able to detect the states, but only mpH$^2$MM is able to determine the $S_{raw}$ values directly.

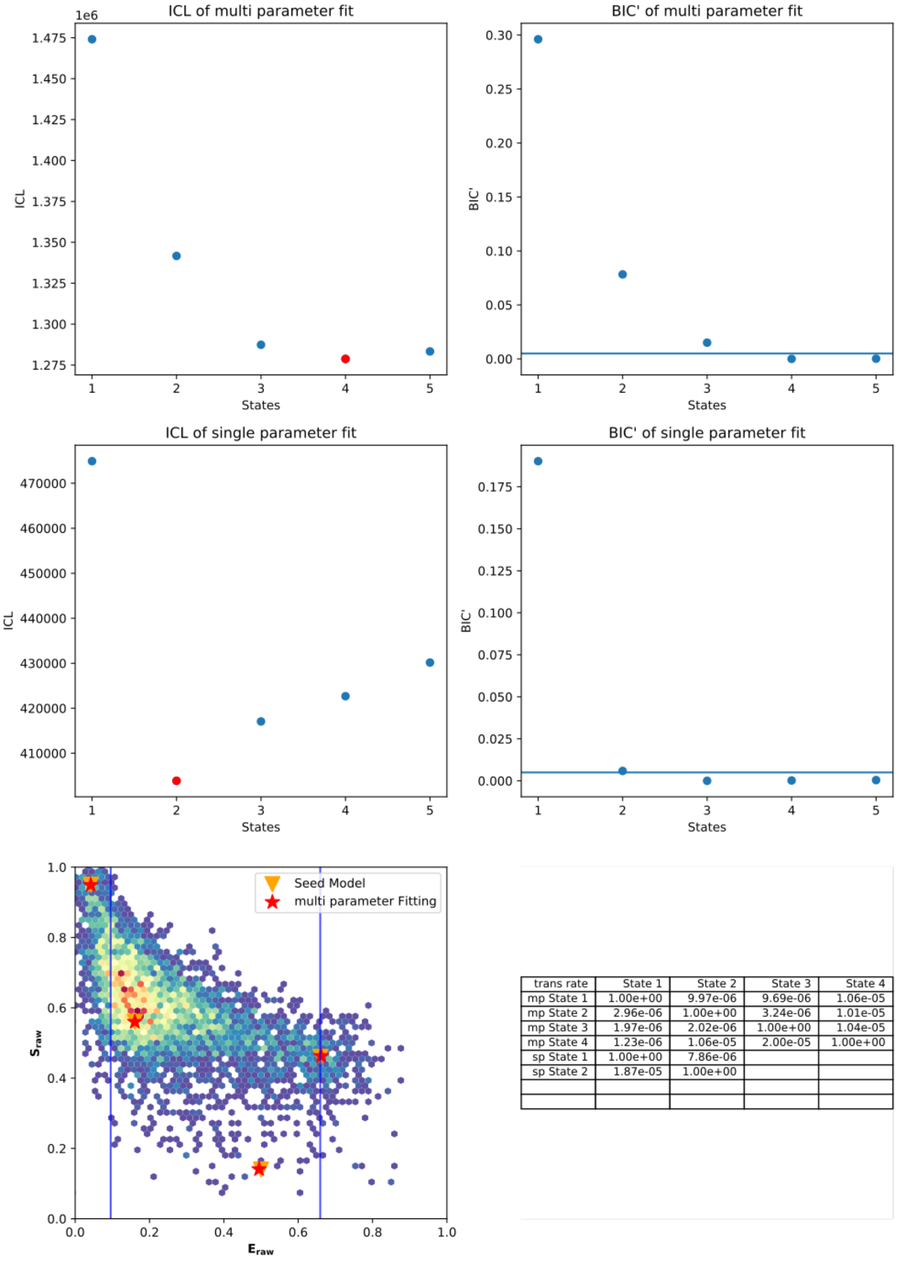

| trans rate | State 1 | State 2 | State 3 | State 4 |
|---|---|---|---|---|
| mp State 1 | 1.00e+00 | 9.97e-06 | 9.69e-06 | 1.06e-05 |
| mp State 2 | 2.96e-06 | 1.00e+00 | 3.24e-06 | 1.01e-05 |
| mp State 3 | 1.97e-06 | 2.02e-06 | 1.00e+00 | 1.04e-05 |
| mp State 4 | 1.23e-06 | 1.06e-05 | 2.00e-05 | 1.00e+00 |
| sp State 1 | 1.00e+00 | 7.86e-06 | | |
| sp State 2 | 1.87e-05 | 1.00e+00 | | |

**figure** S4: **Simulation of a 4 state system with dark donor and dark acceptor state** Note how spH$^2$MM averages the dark acceptor and low FRET states.

##### S5.1.3 HP3 figures

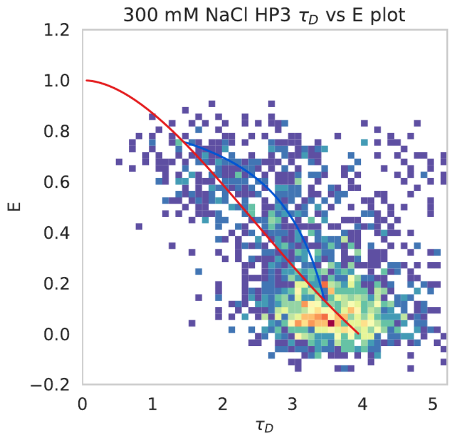

**figure** S5: E-$\tau_D$ of 300 mM NaCl HP3

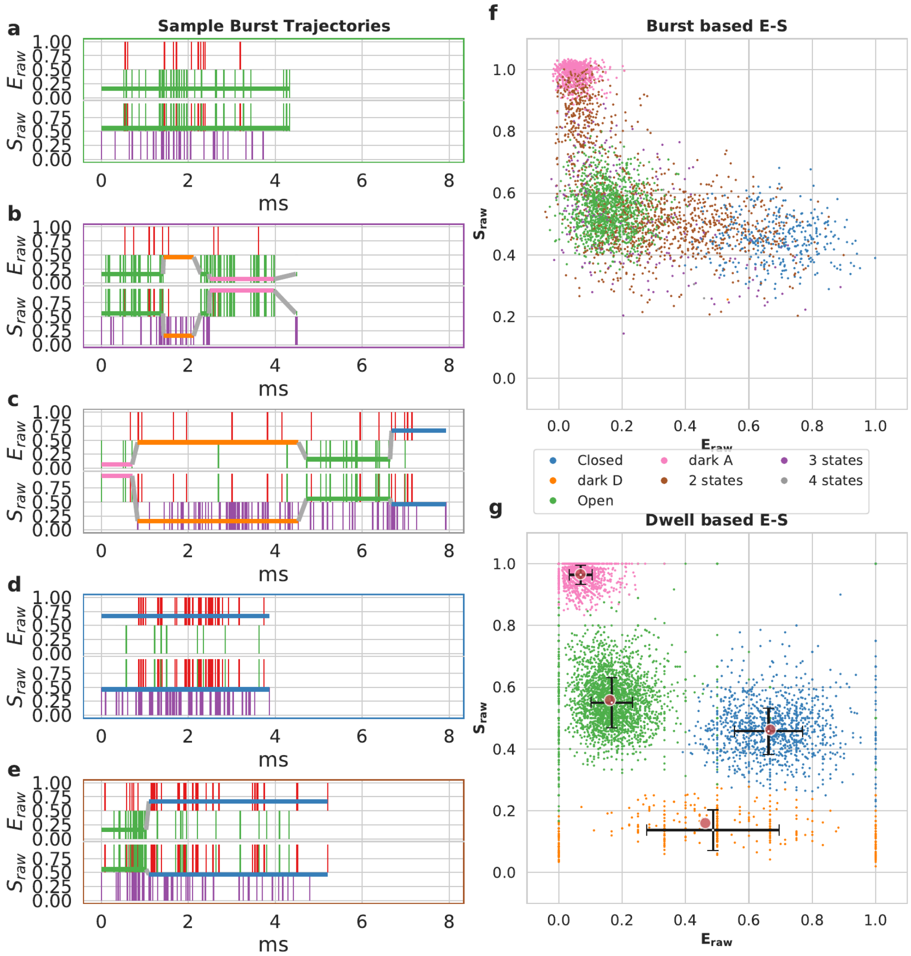

**figure** S6: ***Viterbi* analysis for DNA hairpin at 300 mM NaCl.** a-e) Sample burst trajectories, photons are represented by vertical colored bars, with donor excitation photons colored green or red for donor or acceptor photons, respectively, and purple for acceptor photons arising during acceptor excitation. The most likely state-path determined by the *Viterbi* algorithm is shown by horizontal lines, with upper panel showing the $E_{raw}$ , and the lower panel showing the $S_{raw}$. The color of each segment serves to identify the state, each segment is referred to as a dwell. The color of the border around the trajectory indicates which state(s) are present in the entire burst. f,g) E-S plots of f) bursts, and g) dwells, with colors identical to the borders and dwells in a-e). Red dots in g) represent the states predicted from the $H^2$MM algorithm, and black crosses represent the standard deviations of $E_{raw}$ and $S_{raw}$ values of dwells of a given state.

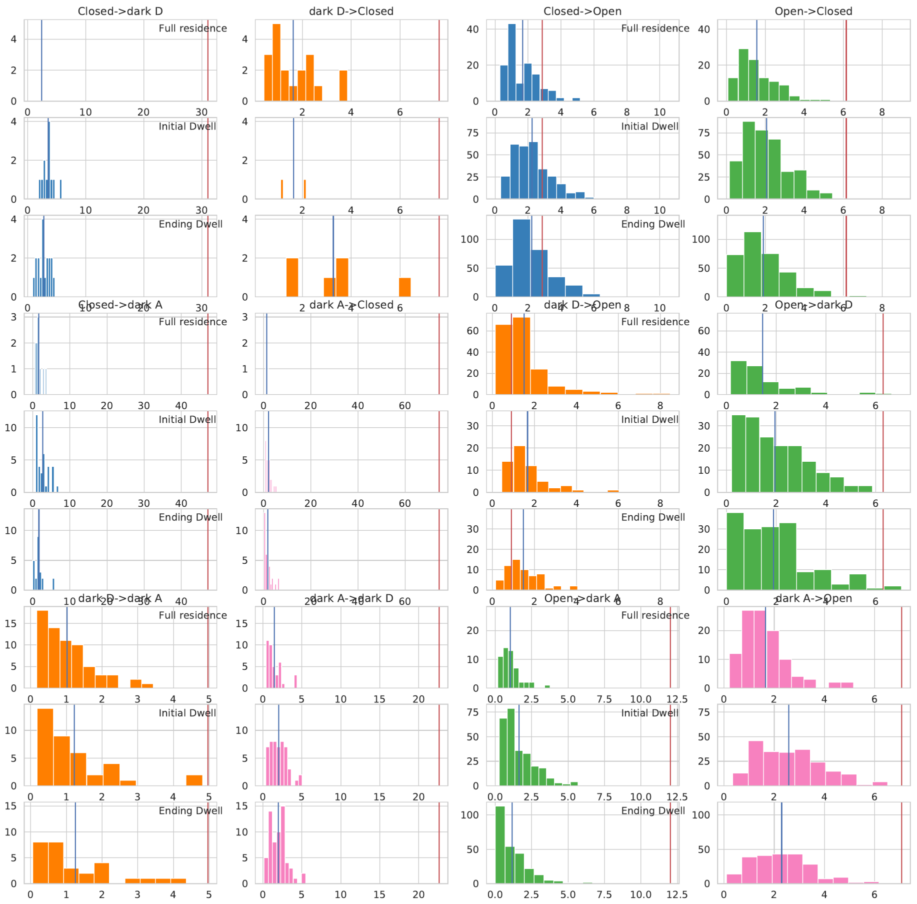

**figure S7: Dwell histograms for DNA hairpin at 300 mM NaCl**

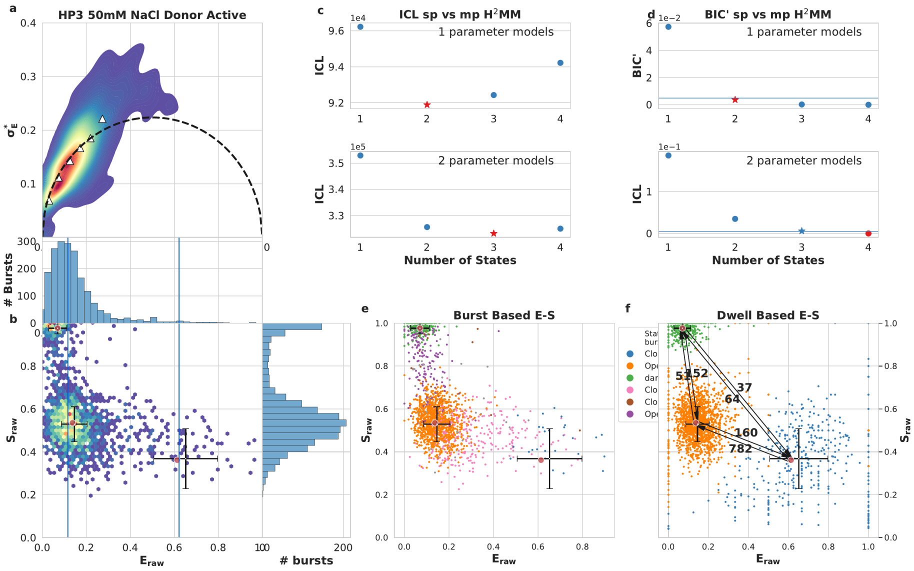

figure S8: **DNA hairpin, 50 mM NaCl** transition rates in s$^{-1}$

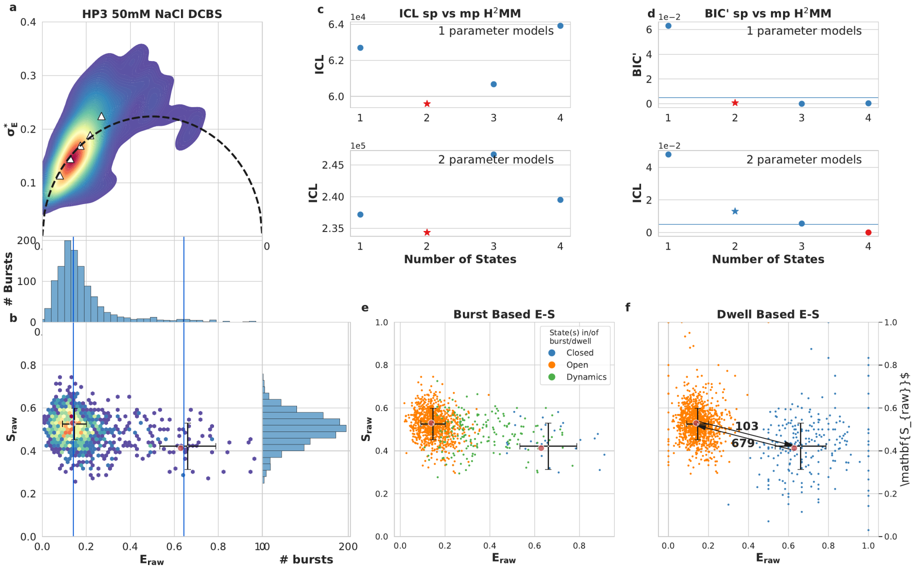

figure S9: **DNA hairpin, 50 mM NaCl** transition rates in s$^{-1}$ using DCBS to remove dark donor/dark acceptor bursts

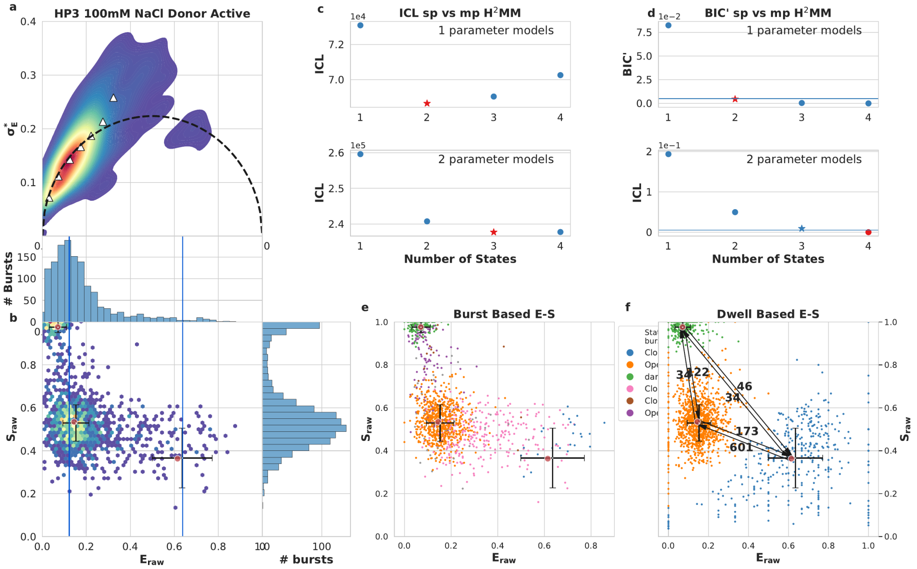

**figure** S10: **DNA hairpin, 100 mM NaCl** transition rates in s$^{-1}$

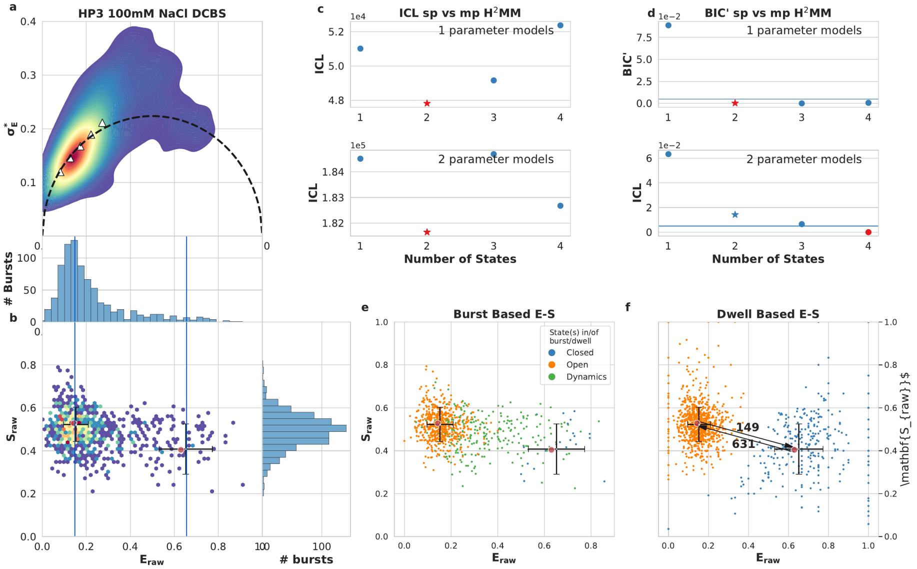

**figure** S11: **DNA hairpin, 100 mM NaCl** transition rates in s$^{-1}$ using DCBS to remove dark donor/dark acceptor bursts

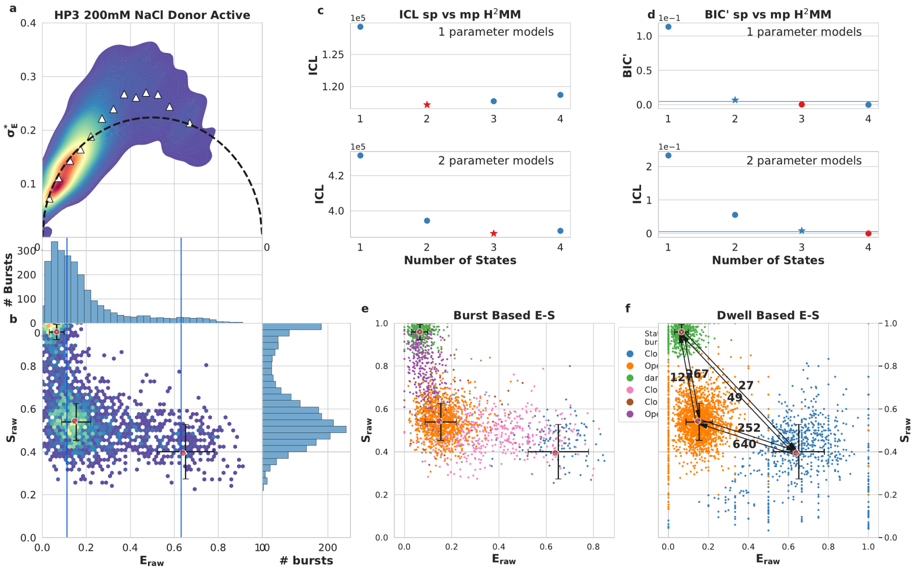

**figure** S12: **DNA hairpin, 200 mM NaCl** transition rates in s$^{-1}$

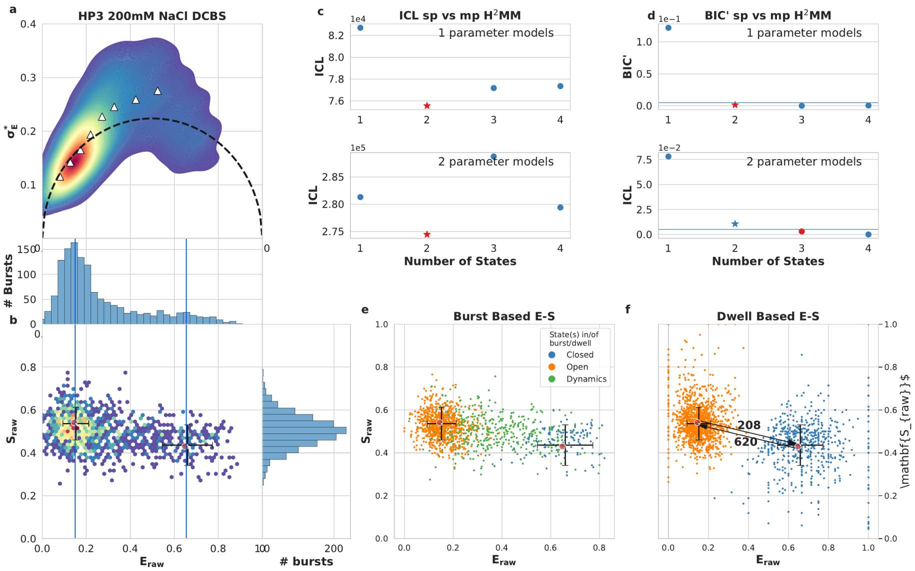

**figure** S13: **DNA hairpin, 200 mM NaCl** transition rates in s$^{-1}$ using DCBS to remove dark donor/dark acceptor bursts

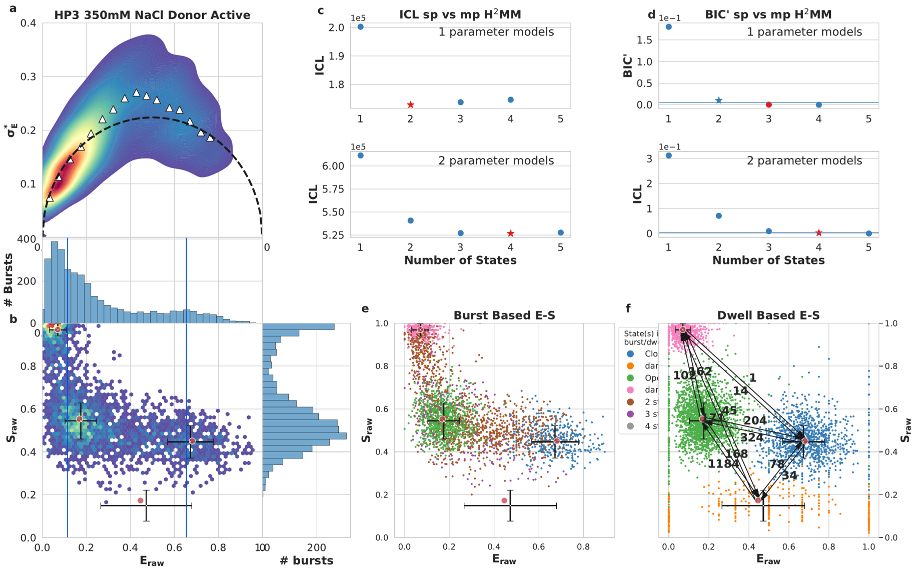

figure S14: **DNA hairpin, 350 mM NaCl** transition rates in s$^{-1}$

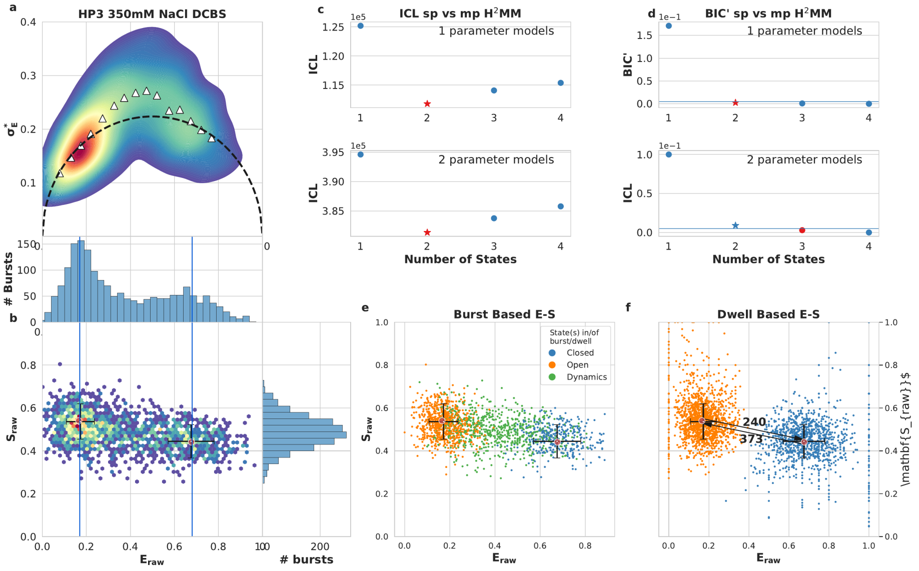

figure S15: **DNA hairpin, 350 mM NaCl** transition rates in s$^{-1}$ using DCBS to remove dark donor/dark acceptor bursts

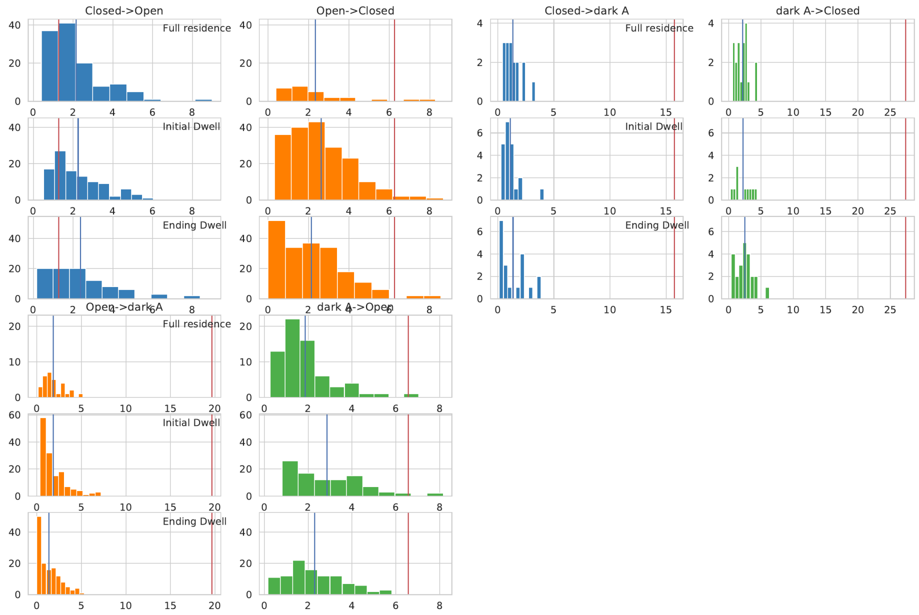

**figure** S16: **Dwell histograms for DNA hairpin at 50 mM NaCl**

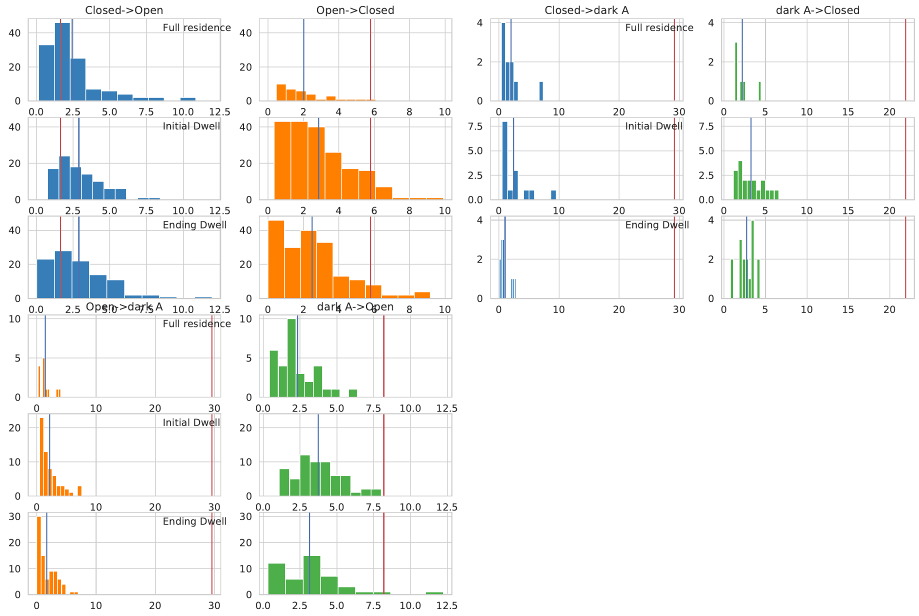

**figure** S17: **Dwell histograms for DNA hairpin at 100 mM NaCl**

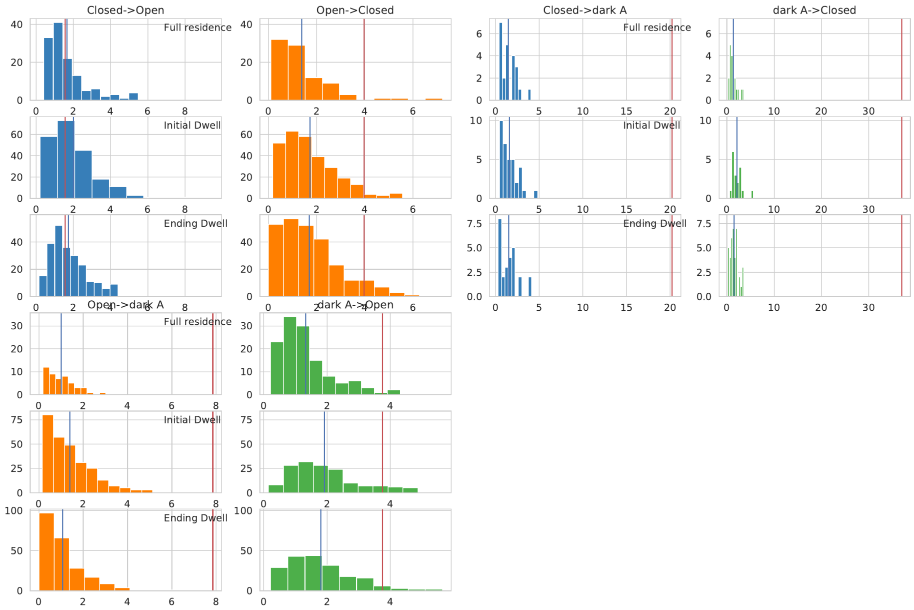

**figure** S18: **Dwell histograms for DNA hairpin at 200 mM NaCl**

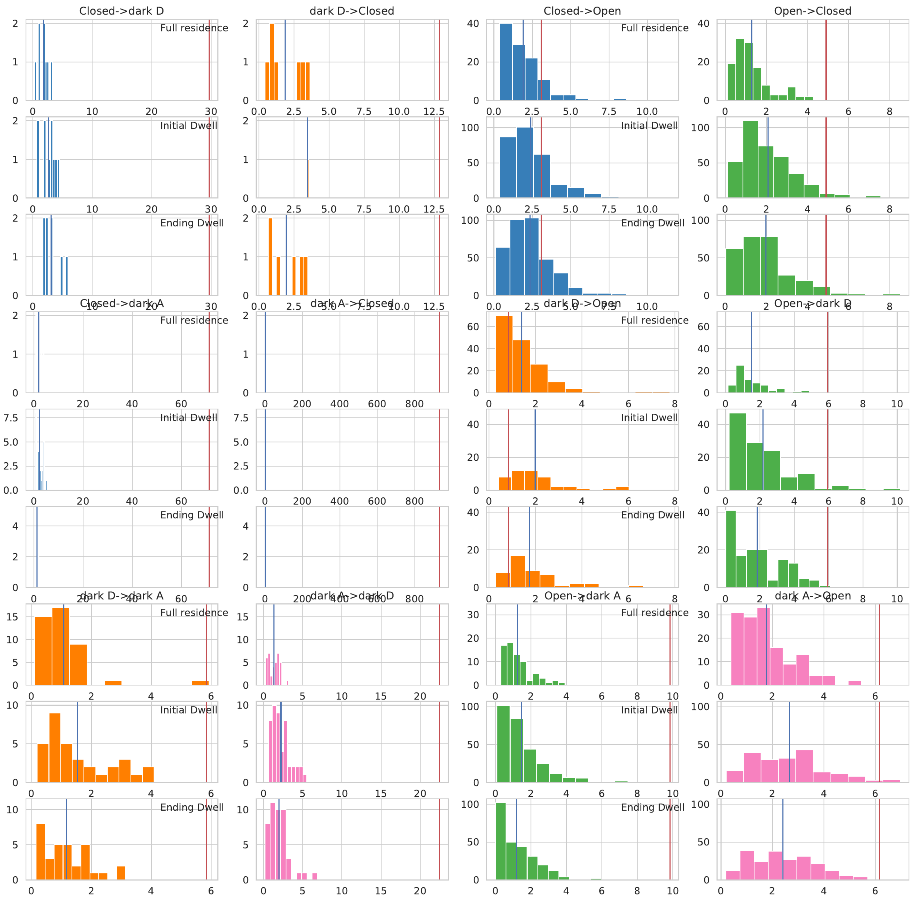

**figure** S19: **Dwell histograms for DNA hairpin at 350 mM NaCl**

##### S5.1.4 MalE

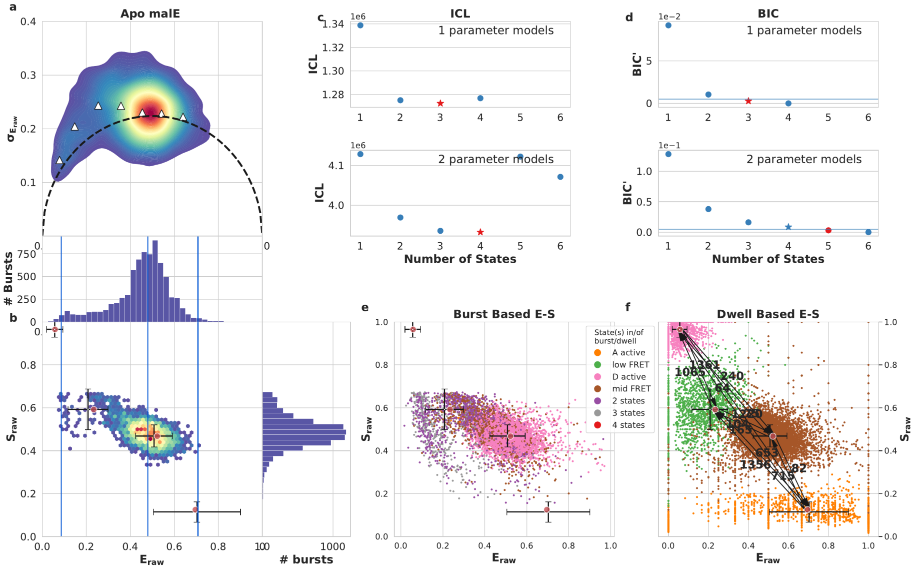

figure S20: **Apo MalE** transition rates in $s^{-1}$

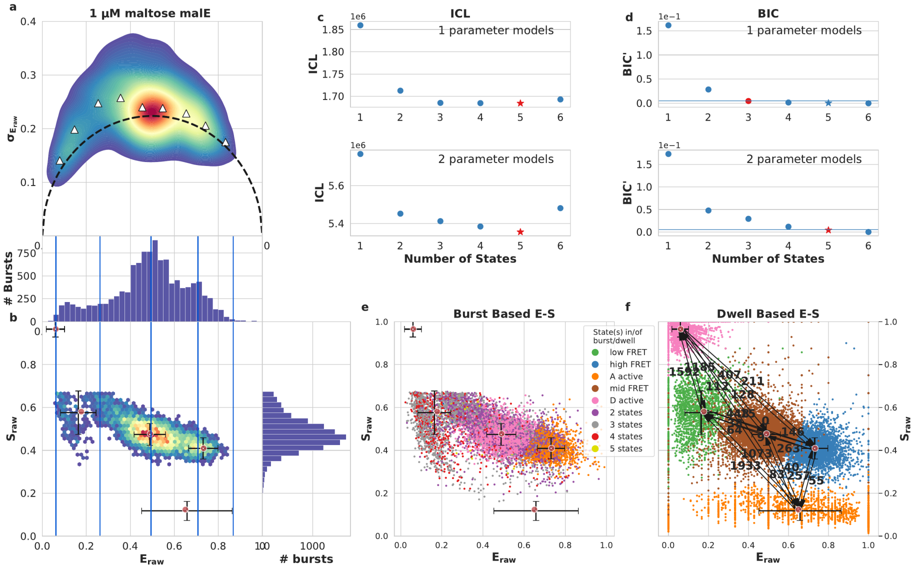

figure S21: **1** μM **maltose MalE** transition rates in $s^{-1}$

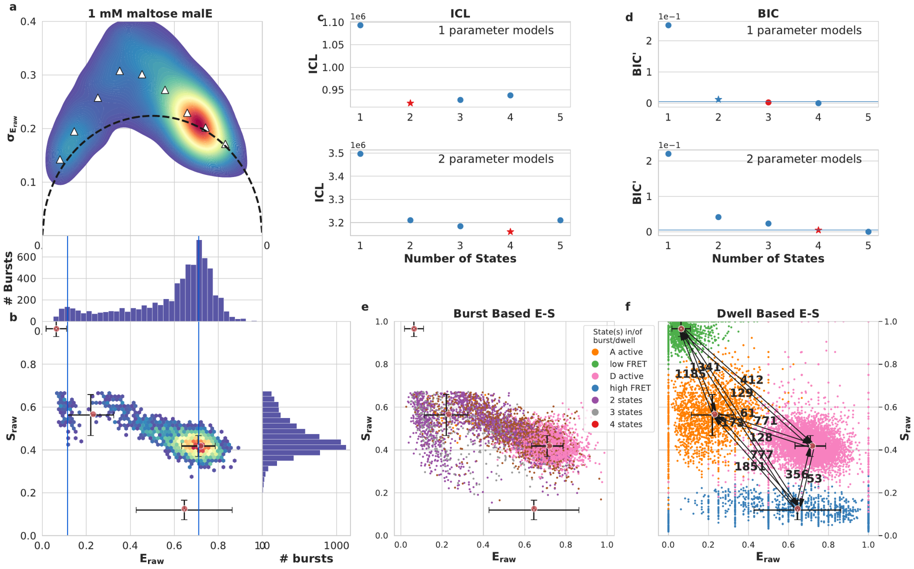

figure S22: **1** mM **maltose MalE** transition rates in $s^{-1}$

###### S5.1.5 YopO

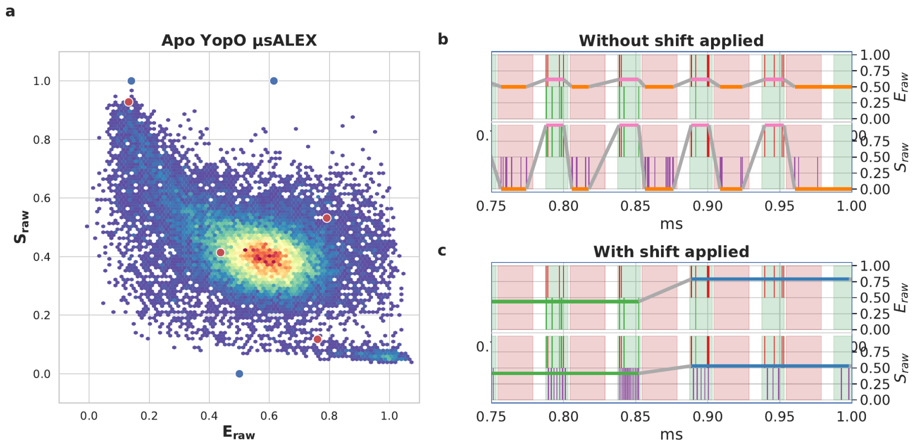

**figure** S23: **Means to adapt mpH$^2$MM to µsALEX measurement** a) E-S plot of apo YopO bursts, with mpH$^2$MM optimized model values overlayed as blue dots for optimization without the µsALEX fix, and red with the µsALEX fix. Note how without the fix, all $S_{raw}$ values are either 0 or 1. b,c) graphical representation of µsALEX fix. A portion of a burst with acceptor excitation photons shifted. Shading indicates excitation period, green and red for donor and acceptor excitation, respectively. Donor and acceptor photons are represented as vertical colored bars. The upper panels contains only donor emission photons, green and red for donor and acceptor emission photons, respectively, while the lower panels adds purple photons for acceptor excitation, acceptor emission photons. The horizontal line indicates the *Viterbi* most likely path, with the $E_{raw}$ value represented in the upper panels, and the $S_{raw}$ value represented in the lower panels. a) Depicts a raw burst trajectory, which results in mpH$^2$MM detecting the alternation period instead of real states. b) Shows a solution to this problem, by taking all acceptor photons in a given acceptor excitation period, and distributing them evenly in the preceding donor excitation period, the alternation period is no longer detected, but mpH$^2$MM will not be able to detect changes in $S_{raw}$ at rates faster than the alternation period.

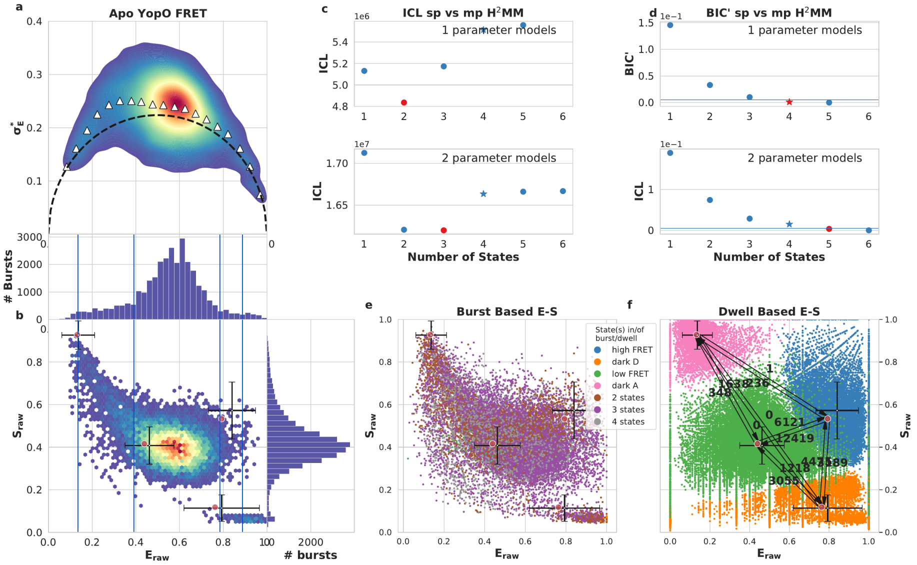

figure S24: **apo YopO** transition rates in $s^{-1}$, 50 µs alternation period

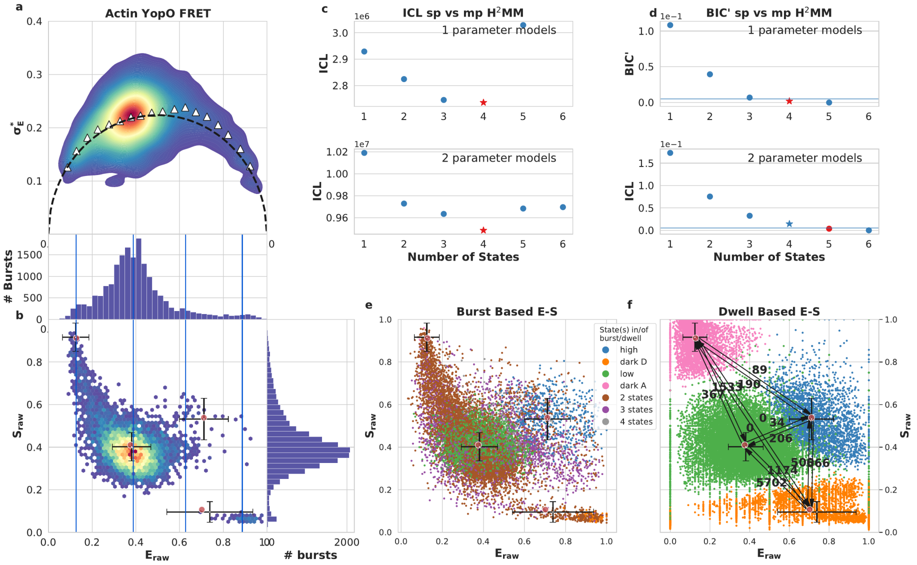

figure S25: **60** µM **actin YopO** transition rates in $s^{-1}$, 50 µs alternation period

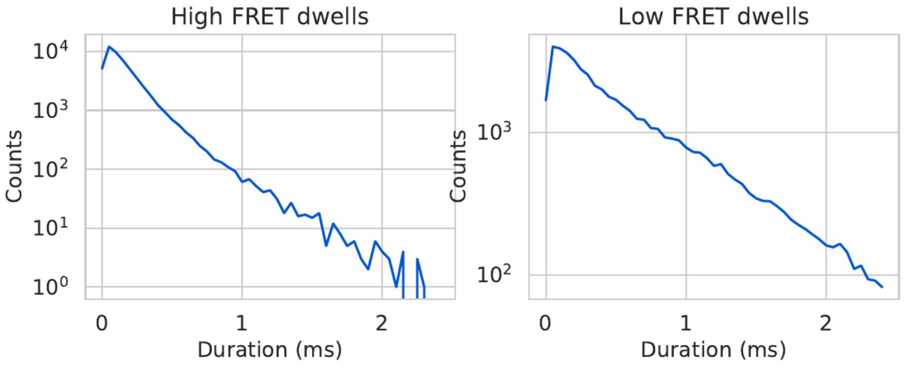

**figure** S26: **Dwell time histograms for low to high and high to low FRET transition rates of YopO** 50 µs alternation period data, demonstrating that dwells are majorly distributed exponentially.

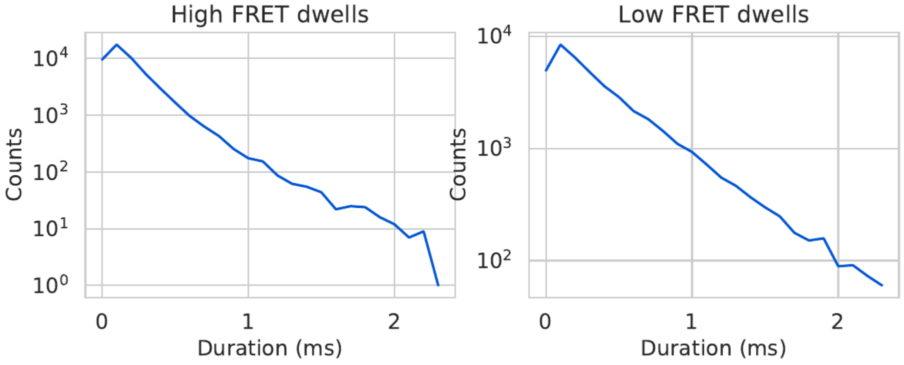

**figure** S27: **Dwell time histograms for low to high and high to low FRET transition rates of YopO** 100 µs alternation period data, demonstrating that wells are majorly distributed exponentially.

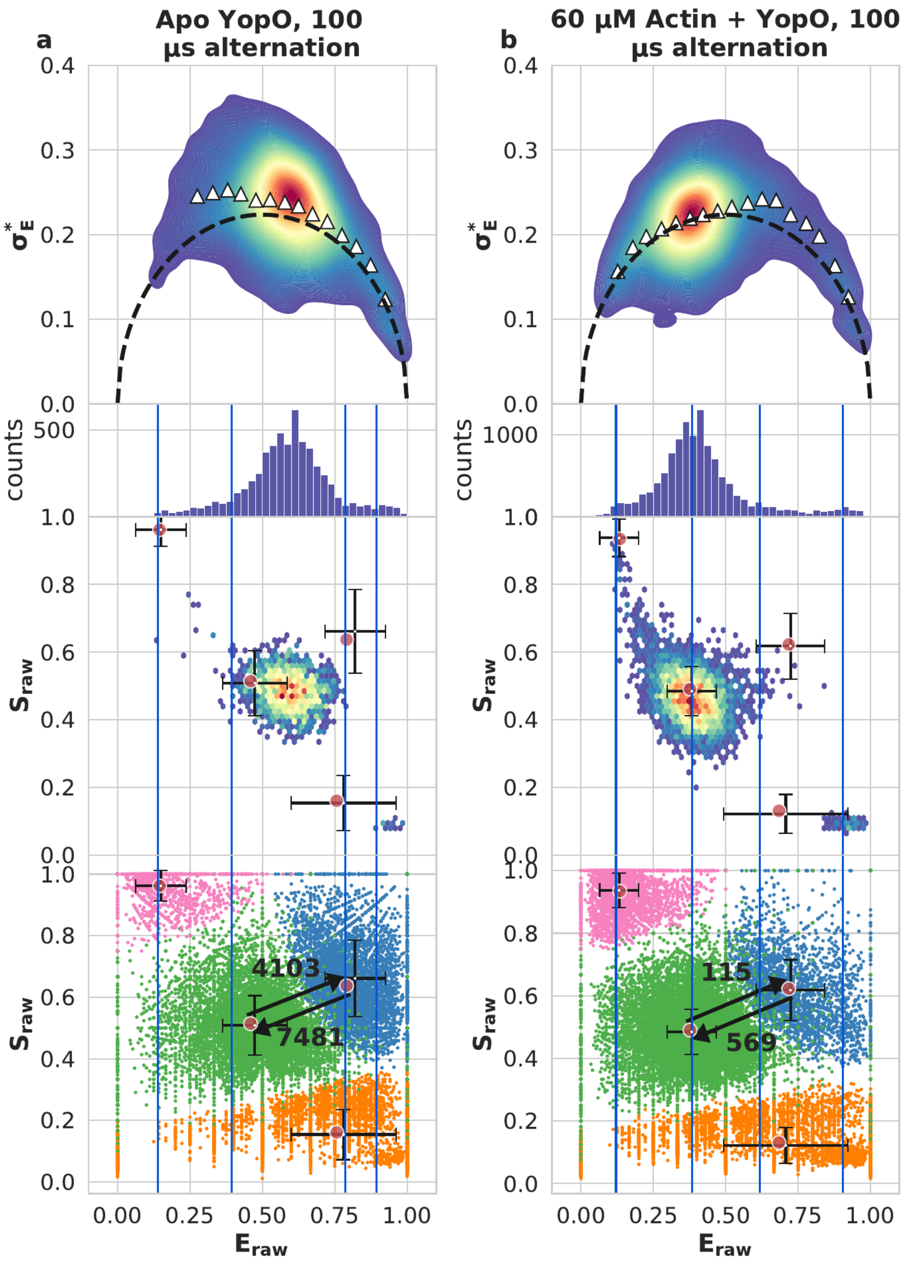

**figure** S28: YopO H$^2$MM analysis with 100 µs alternation period

#### S5.2 Tables

**Table** S1: **Ground truth parameters for simulations (figures S3, S4)** Units of transition rates are the inverse clock period of the simulation. Photons arrive with exponentially distributed delays, with a mean of 1100 clock period. This approximates a clock period of 50 ns in nsALEX experiments.

| Simulation of 3 states with similar $S_{raw}$ (figure S3) | | | | | | |
|---|---|---|---|---|---|---|
| | to State 1 | to State 2 | to State 3 | – | E | S |
| State 1 | – | 1e-6 | 1e-6 | | 0.75 | 0.4 |
| State 2 | 1e-6 | – | 1e-6 | | 0.6 | 0.5 |
| State 3 | 1e-6 | 1e-6 | – | | 0.33 | 0.5 |

| Simulation of 4 states dark donor and dark acceptor state (figure S4) | | | | | | |
|---|---|---|---|---|---|---|
| | to State 1 | to State 2 | to State 3 | to State 4 | E | S |
| State 1 | – | 1e-5 | 1e-6 | | 0.042 | 0.95 |
| State 2 | 2e-5 | – | 2e-6 | 2e-6 | 0.161 | 0.56 |
| State 3 | 2e-5 | 3e-6 | – | 3e-6 | 0.66 | 0.46 |
| State 4 | 1e-5 | 1e-5 | 1e-5 | – | 0.5 | 0.16 |

**Table** S2: $E_{raw}$ and $S_{raw}$ values for DNA hairpin at 300 mM NaCl

| | spH$^2$MM | | mpH$^2$MM | |
|---|---|---|---|---|
| | 2-state | 3-state | 4-state | |
| ICL[a] | 275158.975068564 | 275353.375540217 | 756092.749982401 | |
| | $E_{raw}$ | $E_{raw}$ | $E_{raw}$ | $S_{raw}$ |
| Open[b] | 0.09 | 0.18 | 0.16 | 0.56 |
| Closed[c] | 0.63 | 0.68 | 0.67 | 0.46 |
| dark A[d] | – | 0.07 | 0.07 | 0.97 |
| dark D[e] | – | – | 0.46 | 0.17 |

[a]ICL values are not directly comparable between spH$^2$MM and mpH$^2$MM, as the data set treated by spH$^2$MM probes less photons than by mpH$^2$MM.

[b]The state whose $E_{raw}$, and $S_{raw}$ values best correspond to that predicted for the open conformation, namely a $E_{raw}$≈0.17, and $S_{raw}$≈0.5

[c]The state whose $E_{raw}$, and $S_{raw}$ values best correspond to that predicted for the open conformation, namely a $E_{raw}$≈0.65, and $S_{raw}$≈0.5

[d]Dark Acceptor, identified in the three-state spH$^2$MM as the state with the lowest $E_{raw}$, while more easily identified as having a $E_{raw}$≈0 and $S_{raw}$≈1

[e]Dark Donor, identified as having an $S_{raw}$≈0, not detectable in spH$^2$MM

**Table** S3: The transition rate constants for selected $H^2MM$ models of the DNA hairpin at 300 mM NaCl, states are named as in table S2

| Open & Closed all values in $s^{-1}$ | | | | | |
|---|---|---|---|---|---|
| Open to Closed | | | Closed to Open | | |
| 2-state sp[a] | 3-state sp[b] | 4-state mp[c] | 2 state sp | 3 state sp | 4 state mp |
| 90 | 217 | 157 | 407 | 409 | 349 |
| Open & dark A | | | | | |
| Open to dark A | | | dark A to Open | | |
| 2 state sp | 3 state sp | 4 state mp | 2 state sp | 3 state sp | 4 state mp |
| – | *20*[d] | 99 | – | *92*[d] | 98 |
| Closed & dark A | | | | | |
| Closed to dark A | | | dark A to Closed | | |
| 2 state sp | 3 state sp | 4 state mp | 2 state sp | 3 state sp | 4 state mp |
| – | *57*[d] | 18 | – | *3*[d] | 12 |
| Open & dark D | | | | | |
| Open to dark D | | | dark D to Open | | |
| 2 state sp | 3 state sp | 4 state mp | 2 state sp | 3 state sp | 4 state mp |
| – | – | 168 | – | – | 1043 |
| Closed & dark D | | | | | |
| Closed to dark D | | | dark D to Closed | | |
| 2 state sp | 3 state sp | 4 state mp | 2 state sp | 3 state sp | 4 state mp |
| – | – | *28*[d] | – | – | 135 |
| dark A & dark D | | | | | |
| dark A to dark D | | | dark D to dark A | | |
| 2 state sp | 3 state sp | 4 state mp | 2 state sp | 3 state sp | 4 state mp |
| – | – | 46 | – | – | 726 |

[a] 2-state sp$H^2MM$
[b] 3-state sp$H^2MM$
[c] 4-state mp$H^2MM$
[d] Too few such dwells in all bursts (less than 10)

**Table S4: $E_{raw}$ and $S_{raw}$ values of MalE models**

| State/Condition<br>Low FRET (L) | $E_{raw}$ | $S_{raw}$ |
|---|---|---|
| Apo | 0.16 | 0.57 |
| 1 µM maltose | 0.09 | 0.56 |
| 1 mM maltose | 0.16 | 0.54 |
| Mid FRET (M) | | |
| Apo | 0.47 | 0.44 |
| 1 µM maltose | 0.44 | 0.45 |
| 1 mM maltose | | |
| High FRET (H) | | |
| Apo | – | – |
| 1 µM maltose | 0.70 | 0.38 |
| 1 mM maltose | 0.69 | 0.39 |
| Donor Only (DO) | | |
| Apo | 0.01 | 0.96 |
| 1 µM maltose | 0.01 | 0.96 |
| 1 mM maltose | 0.01 | 0.96 |
| Acceptor only (AO) | | |
| Apo | 0.44 | 0.07 |
| 1 µM maltose | 0.35 | 0.07 |
| 1 mM maltose | 0.35 | 0.07 |

**Table** S5: **MalE transition rates** all rates given in $s^{-1}$

| Condition | L to M | M to L | L to H | H to L | M to H | H to M |
|---|---|---|---|---|---|---|
| Apo | 1220 | 165 | – | – | – | – |
| 1 µM maltose | 448 | 64.2 | 0 | 9.42 | 146 | 263 |
| 1 mM maltose | – | – | 771 | 128 | – | – |
| – | DO to L | L to DO | DO to M | M to DO | DO to H | H to DO |
| Apo | 1360 | 1060 | 240 | 63.9 | – | – |
| 1 µM maltose | 1180 | 1590 | 407 | 112 | 221 | 128 |
| 1 mM maltose | 1340 | 1180 | – | – | 412 | 129 |
| – | AO to L | L to AO | AO to M | M to AO | AO to H | H to AA |
| Apo | 1360 | 653 | 715 | 81.6 | – | – |
| 1 µM maltose | 1930 | 1070 | 83.5 | 257 | 54.7 | |
| 1 mM maltose | | | | | | |
| – | Do to AO | AO to DO | – | – | – | – |
| Apo | 75.1 | 104 | – | – | – | – |
| 1 µM maltose | 14.8 | 38.7 | – | – | – | – |
| 1 mM maltose | 60.7 | 173 | – | – | – | – |

**Table** S6: $\boldsymbol{E_{raw}}$ **and** $\boldsymbol{S_{raw}}$ **values of YopO models, 50 µs alternation period**

| State/Condition<br>Low FRET (L) | $E_{raw}$ | $S_{raw}$ |
|---|---|---|
| Apo | 0.44 | 0.42 |
| 60 µM Actin | 0.38 | 0.41 |
| High FRET (H) | | |
| Apo | 0.79 | 0.53 |
| 60 µM Actin | 0.71 | 0.54 |
| Donor Only (DO) | | |
| Apo | 0.13 | 0.93 |
| 60 µM Actin | 0.13 | 0.91 |
| Acceptor only (AO) | | |
| Apo | 0.76 | 0.12 |
| 60 µM Actin | 0.70 | 0.11 |

**Table** S7: **YopO transition rates, 50 µs alternation** all rates given in $s^{-1}$

| Condition | L to H | H to L | DO to L | L to DO | DO to H | H to DO |
|---|---|---|---|---|---|---|
| Apo | 6100 | 12400 | 1670 | 348 | 1.24 | 236 |
| Actin | 33.7 | 206 | 1530 | 367 | 89.4 | 190 |
| | AO to L | L to AO | AO to H | H to AO | DO to AO | AO to DO |
| Apo | 3050 | 1220 | 4460 | 3190 | 0.00 | 0.00 |
| Actin | 5700 | 1170 | 500 | 566 | 0.00 | 0.00 |

**Table** S8: $\boldsymbol{E_{raw}}$ **and** $\boldsymbol{S_{raw}}$ **values of YopO models, 100 µs alternation period**

| State/Condition Low FRET (L) | $E_{raw}$ | $S_{raw}$ |
|---|---|---|
| Apo | 0.44 | 0.51 |
| 60 µM Actin | 0.38 | 0.49 |
| High FRET (H) | | |
| Apo | 0.78 | 0.62 |
| 60 µM Actin | 0.72 | 0.62 |
| Donor Only (DO) | | |
| Apo | 0.14 | 0.93 |
| 60 µM Actin | 0.13 | 0.93 |
| Acceptor only (AO) | | |
| Apo | 0.73 | 0.16 |
| 60 µM Actin | 0.68 | 0.13 |

**Table** S9: **YopO transition rates, 100 µs alternation** all rates given in $s^{-1}$

| Condition | L to H | H to L | DO to L | L to DO | DO to H | H to DO |
|---|---|---|---|---|---|---|
| Apo | 5570 | 100000 | 1440 | 288 | 43.5 | 345 |
| Actin | 115 | 569 | 1270 | 334 | 73.0 | 273 |
| | AO to L | L to AO | AO to H | H to AO | DO to AO | AO to DO |
| Apo | 3710 | 1820 | 3350 | 2200 | 0.00 | 0.00 |
| Actin | 5310 | 1120 | 704 | 848 | 0.00 | 0.00 |

**Table** S10: *Durations of hairpin notebooks*

| Analysis | Time | Type | No. of optimizations |
|---|---|---|---|
| HP3 50 mM NaCl | 28 min 15.06 s | Notebook | 16 |
| HP3 50 mM NaCl DA spH$^2$MM | 10 min 22.48 s | spH$^2$MM | 4 |
| HP3 50 mM NaCl DA mpH$^2$MM | 22.9660 seconds | mpH$^2$MM | 4 |
| HP3 50 mM NaCl DCBS spH$^2$MM | 6 min 57.18 s | spH$^2$MM | 4 |
| HP3 50 mM NaCl DCBS mpH$^2$MM | 9 min 27.30 s | mpH$^2$MM | 4 |
| HP3 100 mM NaCl | 20 min 4.68 s | Notebook | 16 |
| HP3 100 mM NaCl DA spH$^2$MM | 5 min 0.22 s | spH$^2$MM | 4 |
| HP3 100 mM NaCl DA mpH$^2$MM | 16.7651 seconds | mpH$^2$MM | 4 |
| HP3 100 mM NaCl DCBS spH$^2$MM | 5 min 5.79 s | spH$^2$MM | 4 |
| HP3 100 mM NaCl DCBS mpH$^2$MM | 8 min 39.06 s | mpH$^2$MM | 4 |
| HP3 200 mM NaCl | 14 min 9.45 s | Notebook | 16 |
| HP3 200 mM NaCl DA spH$^2$MM | 3 min 52.96 s | spH$^2$MM | 4 |
| HP3 200 mM NaCl DA mpH$^2$MM | 20.4220 seconds | mpH$^2$MM | 4 |
| HP3 200 mM NaCl DCBS spH$^2$MM | 5 min 57.05 s | spH$^2$MM | 4 |
| HP3 200 mM NaCl DCBS mpH$^2$MM | 2 min 50.85 s | mpH$^2$MM | 4 |
| HP3 250 mM NaCl | 23 min 57.20 s | Notebook | 17 |
| HP3 250 mM NaCl DA spH$^2$MM | 3 min 2.25 s | spH$^2$MM | 4 |
| HP3 250 mM NaCl DA mpH$^2$MM | 8 min 31.26 s | mpH$^2$MM | 5 |
| HP3 250 mM NaCl DCBS spH$^2$MM | 8 min 32.98 s | spH$^2$MM | 4 |
| HP3 250 mM NaCl DCBS mpH$^2$MM | 2 min 26.08 s | mpH$^2$MM | 4 |
| HP3 300 mM NaCl | 17 min 52.06 s | Notebook | 17 |
| HP3 300 mM NaCl DA spH$^2$MM | 3 min 12.62 s | spH$^2$MM | 4 |
| HP3 300 mM NaCl DA mpH$^2$MM | 1 min 41.21 s | mpH$^2$MM | 4 |
| HP3 300 mM NaCl DCBS spH$^2$MM | 7 min 50.71 s | spH$^2$MM | 4 |
| HP3 300 mM NaCl DCBS mpH$^2$MM | 1 min 5.34 s | mpH$^2$MM | 4 |
| HP3 350 mM NaCl | 19 min 12.88 s | Notebook | 17 |
| HP3 350 mM NaCl DA spH$^2$MM | 6 m 2.99 s | spH$^2$MM | 4 |
| HP3 350 mM NaCl DA mpH$^2$MM | 1 min 46.51 s | mpH$^2$MM | 5 |
| HP3 350 mM NaCl DCBS spH$^2$MM | 7 min 20.54 s | spH$^2$MM | 4 |
| HP3 350 mM NaCl DCBS mpH$^2$MM | 2 min 43.13 s | mpH$^2$MM | 4 |

**Table** S11: **Duration of MalE and YopO notebooks**

| Analysis | Time | Type | No. of optimizations |
|---|---|---|---|
| MalE notebook | 4 hr 56 min 21.03 s | Notebook | 31 |
| MalE apo spH$^2$MM | 6 min 14.09 s | spH$^2$MM | 4 |
| MalE apo mpH$^2$MM | 1 hr 42 min 33.61 s | mpH$^2$MM | 6 |
| MalE 1 µM maltose spH$^2$MM | 51 min 47.01 s | spH$^2$MM | 6 |
| MalE 1 µM maltose mpH$^2$MM | 1 hr 32 min 9.53 s | mpH$^2$MM | 6 |
| MalE 1 mM maltose spH$^2$MM | 3 min 11.88 s | spH$^2$MM | 4 |
| MalE 1 mM maltose mpH$^2$MM | 23 min 38.60 s | mpH$^2$MM | 5 |
| YopO 50 µs alternation notebook | 9 hr 22 min 48.33 s | Notebook | 17 |
| YopO Apo spH$^2$MM | 1 hr 48 min 16.04 s | spH$_2$MM | 5 |
| YopO Apo mpH$^2$MM | 3 hr 31 min 0.29 s | mpH$^2$MM | 6 |
| YopO actin spH$^2$MM | 45 min 36.22 s | mpH$^2$MM | 6 |
| YopO actin mpH$^2$MM | 1 hr 51 min 45.84 s | mpH$^2$MM | 6 |
| YopO 100 µs alternation notebook | 6hr 18 min 7.69 s | Notebook | 17 |
| YopO Apo spH$^2$MM | 1 hr 52 min 52.20 s | spH$_2$MM | 5 |
| YopO Apo mpH$^2$MM | 3 hr 19 min 53.64 s | mpH$^2$MM | 6 |
| YopO actin spH$^2$MM | 30 min 42.07 s | mpH$^2$MM | 6 |
| YopO actin mpH$^2$MM | 30 min 5.47 s | mpH$^2$MM | 6 |

1.5004606. URL: http://dx.doi.org/10.1063/1.5004606%20http://aip.scitation.org/doi/10.1063/1.5004606.
